# Differentiating unirradiated mice from those exposed to conventional or FLASH radiotherapy using MRI

**DOI:** 10.1101/2025.02.01.636061

**Authors:** Jeannette Jansen, Adam Kimbler, Olivia Drayson, Bernard Lanz, Jessie Mosso, Veljko Grilj, Benoit Petit, Javier Franco-Perez, Aaron Simon, Charles L. Limoli, Marie-Catherine Vozenin, Craig Stark, Paola Ballesteros-Zebadua

## Abstract

**Background and purpose:** The FLASH effect expands the therapeutic ratio of tumor control to normal tissue toxicity observed after delivery of ultra-high (>100 Gy/s FLASH-RT) vs. conventional dose rate radiation (CONV-RT). In this first exploratory study, we assessed whether ex-vivo Magnetic Resonance Imaging (MRI) could reveal long-term differences after FLASH-RT and CONV-RT whole-brain irradiation.

**Materials and methods:** Female C57BL/6 mice were divided into three groups: control (non-irradiated), conventional (CONV-RT 0.1 Gy/s), and ultra-high dose rates (FLASH-RT 1 pulse, 5.5 x 10˄6 Gy/s), and received 10 Gy of whole-brain irradiation in a single fraction at 10 weeks of age. Mice were evaluated by Novel Object Recognition cognitive testing at 10 months post-irradiation and were sampled at 13 months post-irradiation. Ex-vivo brains were imaged with a 14.1 Tesla/26 cm magnet with a multimodal MRI protocol, including T2-weighted TurboRare (T2W) and diffusion-weighted imaging (DWI) sequences.

**Results:** In accordance with previous results, cognitive tests indicated that animals receiving CONV-RT exhibited a decline in cognitive function, while FLASH-RT performed similarly to the controls. MRI showed decreased hippocampal mean intensity in the CONV-RT mice compared to controls but not in the FLASH-RT group. Comparing CONV-RT to control, we found significant changes in multiple whole-brain diffusion metrics, including the mean Apparent Diffusion Coefficient (ADC) and Mean Apparent Propagator (MAP) metrics. By contrast, no significant diffusion changes were found between the FLASH-RT and control groups. In an exploratory analysis compared to controls, regional diffusion metrics were primarily altered in the basal forebrain and the insular cortex after CONV-RT, and after FLASH-RT, a trend reduction was also observed.

**Conclusion:** This study presents initial evidence that MRI can uncover clear changes in the brain after CONV-RT but not after FLASH-RT. The MRI results aligned with the observed cognitive protection after FLASH-RT, indicating the potential use of MRI to analyze the FLASH response.

## Introduction

FLASH-radiotherapy (FLASH-RT) is a promising technology involving ultra-high dose rates (>100 Gy/s) [1]. Remarkably, FLASH-RT reduces normal tissue toxicities while maintaining tumor kill compared to conventional dose rate radiotherapy. The underlying mechanisms are under investigation; however, in the brain, FLASH-RT preserves synaptic plasticity, neuronal structure and decreases neuroinflammation, reducing radiation-induced cognitive toxicity [2, 3, 4]. It, therefore, has significant potential to alleviate the cognitive toxicities that currently impact the quality of life of patients whose treatment requires cranial radiotherapy.

Magnetic Resonance Imaging (MRI), which provides anatomic, microstructural, and functional information, has demonstrated considerable promise as a clinical tool exploring the impact of radiotherapy [5]. To that end, as imaging measurements for radiation-induced cognitive toxicity have been described, they are being incorporated into large prospective trials related to cognitive toxicity mitigating treatment strategies (e.g. clinical trials NCT04588246, NCT03550391) [6, 7].

Structural MRI volumetry has identified changes in the hippocampus associated with cognitive decline [8, 9, 10]. MRI can also measure the relaxation time of biological tissue, which is the time for protons to return to a state of equilibrium after electromagnetic excitation. The transverse component, known as T2, is sensitive to changes in brain tissue microstructure integrity [11, 12, 13], which can be detected prior to tissue atrophy in Alzheimer’s patients [11] and is associated with aging [12] and mild cognitive impairment [13].

Diffusion-weighted imaging (DWI) is another MRI sequence sensitive to the motion of intra-cellular and extracellular water in brain tissue, revealing microstructural tissue properties that may not be detectable with conventional techniques [14]. DWI can detect pathological processes, such as demyelination and inflammation, associated with the toxic effects of cranial irradiation. It can also examine the changes in the microstructure of grey matter as seen with Neurite Orientation Dispersion and Density Imaging (NODDI) [15]. This makes DWI a suitable tool for investigating brain diseases in preclinical models [15, 16]. Obtaining MRI images sensitive to changes in water diffusion allows the use of statistical models to analyze pathological changes in connectivity [14]. In the last few years, these techniques have had promising results as tools to evaluate for cognitive pathologies like Alzheimer’s Disease [17].

By controlling the hydration of sampled tissues, it is possible to preserve the water diffusion properties in ex-vivo brains. Compared to in-vivo imaging, ex-vivo MRI can provide more stable and clean acquisitions since there are no limitations related to anesthesia, scanning time, or animal motion artifacts and that the shimming conditions can be optimized by placing the brain into fluoropolymer oil, which both lacks MR signal and has magnetic susceptibility similar to that of tissue. Here we show that ex-vivo MRI can reveal structural and microstructural alterations in the brain induced by CONV-RT but no significant modifications were measured after FLASH-RT.

## Materials and Methods

### Whole Brain Irradiation

This study included 28 female black C57BL/6 mice, non-tumor-bearing (10 controls, 9 CONV-RT, and 9 FLASH-RT) irradiated at 10 weeks of age. Cohorts received 0 or 10 Gy of whole-brain irradiation in a single fraction using the Oriatron 6e (eRT6) linear accelerator (5.5 MeV electron beam). The eRT6 has been validated extensively to produce the FLASH effect [1]. The CONV-RT dose rate was set to 0.1 Gy/s, while the ultra-high dose rate/FLASH-RT was delivered in 1 single pulse of 10 Gy and a dose rate of 5.5 x 10˄^6^ Gy/s. Mice were irradiated under isofluorane anesthesia (1.5-2.5%) with air as the carrier gas and using a graphite 17 mm collimator focusing the beam to cover the entire brain as already described [1]. All animal procedures were performed in accordance with international and national animal welfare regulations and in compliance with the protocol ethical cantonal authorization no. **VD3603 (Switzerland).**

### Novel Object Recognition Testing

FLASH-RT has been shown to protect against the neurocognitive deficits caused by CONV-RT over protracted post-irradiation intervals [2]. In the current study, we evaluated the long-term impact of FLASH-RT and CONV-RT on cognition using the novel object recognition (NOR) test performed ten months post-irradiation in six mice per group (Fig. 1). The NOR task involved a sequence of habituation (no objects), familiarization (two distinct objects), and a test phase in which one of the prior objects is switched with a new one [1]. Animals with intact cortical function are expected to spend more time with the novel object, whereas animals with compromised memory will not [18]. Exploration time the mice spent with each object was measured via a camera and analyzed by a NoldusEthovision XT system in order to calculate the Discrimination Index (DI) as the equation below.

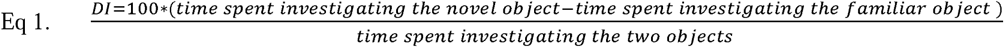

**Figure 1.**
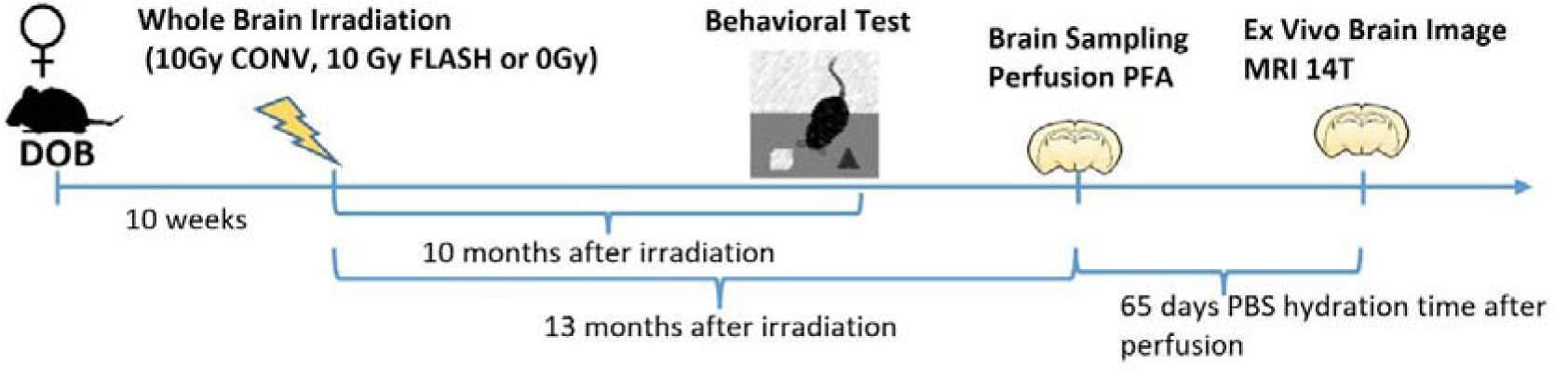
Diagram showing the timeline of performed procedures in female black C57BL/6 mice. All animals were in good health at both the time of the behavioral studies and sampling before the imaging procedures

The Novel Object Recognition test statistical analyses were performed using the Shaphiro-Wilk test to confirm the normality and applying unpaired one-way ANOVA with the Tukey posthoc test. Data points were considered outliers if they deviated by more than 2 standard deviations from the mean and were excluded from the analysis. Results were expressed as mean values ± standard deviations, and the significance level was chosen when p≤.05.

### Sample preparation

13 months post-irradiation mice were intracardiacally perfused with PBS and PFA 4%. The general health status of all the mice was fine at the time of sampling. Brains were extracted, stored in PFA for 48 hrs, and then stored in PBS+azide (0.1 %) for 9.3 weeks (Fig 1.). Ex-vivo MRI imaging was run without any contrast agent. Brains were scanned in pairs inside a plastic holder filled with Fomblin (Fomblin Profludropolyether; Ausimont, Thorofare, NJ). All air bubbles were removed using a vacuum pump for 45 mins.

### Magnetic Resonance Imaging

A 14.1 Tesla/26 cm magnet (Magnex Scientific, Oxford, UK / Bruker Avance Neo, Paravision 360 v1.1, Ettlingen, Germany) was used for ex-vivo imaging equipped with a 12 cm internal diameter gradient coil insert (1000 mT/m, 270 μs rise time) that is well-suited for small-animal imaging. A custom-made transmit-receive RF saddle coil of 24 mm inner diameter was used. Note that 14.1 T is about 5-10 times stronger, and the 1000 mT/m gradients are about 5-20 times stronger than those used in clinical human applications. This MR imaging signal level provides higher spatial resolution and improved diffusion contrasts in small-animal imaging, where small voxel sizes are required. We acquired structural and diffusion-weighted sequences for each pair of brains.

**B0 Map:** B0 map of brains was acquired with two averages, and image size of 96×96×96 and a field of view (FOV) of 34×34×34 mm^3^. This map served for B0 shimming with the Bruker MapShim approach on a cuboid surrounding the brain pairs.

- **T2w TurboRare:** 2D T2 weighted TurboRare structural scans were acquired with an echo time (TE) of 9ms and a repetition time (TR) of 3000 ms, 55 slices and 3 averages, with slice thickness 0.3 mm and slice gap 0.1 mm, slice orientation was axial, and read direction was left-right, image size of 107×150 and a FOV of 15×21 mm^2^.
- **Diffusion Weight Imaging DWI:** Diffusion tensor images were acquired using a Spin-Echo diffusion preparation module and a segmented 2D multislice EPI imaging readout (8 segments). Two scans were acquired with b-values = 2500 and 5000 s/mm² respectively with TE/TR = 28/2000ms, 45 directions, 25 averages, slice thickness 0.6 mm, 40 slices, 0 mm gap, 0.14 mm² resolution, FOV 24×15mm^2^, delta/DELTA: 5/14ms and read orientation ventral-dorsal, to keep the shortest dimension for phase encoding. For image correction purposes, a phase-reversed image was additionally acquired with b = 1000 s/mm². The multi-shell diffusion data was converted and saved for analysis using different metrics. In addition, an A_0_ image was acquired with identical acquisition parameters and b-value = 0 as morphological reference image for tensor data.

### Structural MRI Analysis

Structural image analysis was conducted using the Advanced Normalization Tools (ANTS) [19, 20] and FMRIB Software Library (FSL) [21]. The T2w images were bias-field corrected using ANTS function ‘N4BiasFieldCorrection’ and then registered to the average template space using ANTS ‘antsRegistrationSyn’. A hippocampus mask was created in the average template space using the Allen mouse brain atlas [22]. The hippocampus mask was then transformed to the space of each corrected T2w image using the generic affine and the inverse warp transformations generated during registration with the ANTS ‘antsApplyTransforms’ function with the multi-label interpolator (see Figure 3A). The volume in mm^3^ and mean intensity were calculated from the masked volumes and the whole brains in the T2w images using FSL function ‘fslstats’. In order to calculate the mean intensity across the brain, all images were thresholded using FSL ‘fslmaths’ to remove voxels outside of the brain volume (threshold of 3,000-40,000) and then the cerebellum was removed using FSL ‘fslroi’ due to strong intensity heterogeneity in the posterior part of the brain (see Figure 3A). The mean intensities of the masked volumes were then adjusted with the equation below:

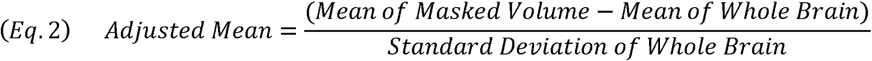

Following volume and intensity calculation, statistical analyses were carried out using GraphPad Prism (version 10) software. All data were first confirmed for normality using the Shapiro-Wilk test. Volumetric and adjusted intensity data were analyzed using one-way ANOVA followed by the Bonferroni multiple comparison test leading to our final, corrected alpha. Data points were considered outliers if they deviated by more than 2 standard deviations from the mean and were excluded from the analysis. Data in the text are presented as means ± standard error, and all analyses considered a value of p ≤ 0.05 to be statistically significant.

### Diffusion MRI Processing

Three mice from each group had shorter PBS hydration times, so they were excluded from the diffusion analysis. One additional mouse in the control group was excluded from the diffusion analysis due to poor quality of the diffusion image scan. Only six mice per treatment group and five in the control were included in the diffusion analysis. Diffusion data was preprocessed using a custom MRTrix-based pipeline [23]. Thermal noise was removed using MRTrix’s ‘dwidenoise’ tool [24]. MRTrix’s ‘mrdegibbs’ was used for Gibbs artifact removal [25]. Eddy current correction was then conducted using MRTrix’s ‘dwifslpreproc’ tool, which wraps FSL’s Eddy tool for eddy current [26], motion [27], susceptibility-induced distortion [28] and bias field correction [29]. After correction for motion and eddy currents, B1-field inhomogeneity was corrected using MRTrix’s ‘dwibiascorrect’ [30] b=0 images were extracted and averaged for use in the registration of diffusion to structural images.

### Calculation of Diffusion Metrics

Diffusion tensor calculations were conducted using MRTrix [23]. Mean apparent diffusion coefficients (ADC) were calculated using ‘dwi2adc’. Tensor metrics were first calculated using ‘dwi2tensor’, and then individual metrics were exported using ‘tensor2metric’. The exported metrics used in our analyses were axial diffusivity (axial diffusivity, same as principal eigenvalue), fractional anisotropy (FA), and radial diffusivity (RD, equal to the mean of the two non-principal eigenvalues).

Neurite orientation dispersion and density imaging (NODDI) metrics were calculated using Microstructure Diffusion Toolbox (MDT) [31] and Accelerated Microstructure Imaging via Convex Optimization (AMICO) [32]. From MDT, Orientation Dispersion Index (ODI) and Neurite Dispersion Index (NDI) were used. From AMICO, equivalent metrics were used: OD for ODI, Intracellular Volume Fraction (ICVF) for NDI) in addition to isotropic volume fraction (ISOVF).

Mean Apparent Propagator (MAP) metrics [33] were calculated using ‘dipy’ (1.5.0) [34]. From ‘dipy’ we get multiple different metrics: the norm of the Laplacian signal (lapnorm), the Q-space Inverse Variance (qiv), the Mean Squared Displacement (msd), the return to the axis probability (rtap), the return to the origin probability (rtop), and the return to plane probability (rtpp).

After calculation in subject space, brain maps containing each metric were subsequently warped from individual space to the Allen Mouse Brain Atlas Space for calculation and group comparisons.

### Diffusion Statistical Analysis

Our initial whole-brain DWI analysis was designed to determine whether the various diffusion metrics differed across groups without any clear *a priori* constraints about where such effects might be observed. With modest sample sizes and with unknown inherent blur (spatial correlation) across adjacent voxels, approaches that would let us clearly isolate specific regions as having reliable differences that would survive multiple comparison corrections were deemed less viable. Instead, using the whole brain as the target region, we opted for an approach that would identify potential differences across treatments agnostic to where they might occur in our primary analysis. By sacrificing the ability to localize the findings, we avoid the limitations imposed by the associated multiple comparison corrections. As a follow-up exploratory analysis, this process was repeated across anatomical ROIs to gain potential insight into possible regional variation or localization of effects.

This analysis began by using AFNI’s 3dANOVA [35, 36] to conduct a voxel-wise one-way ANOVA for each metric to identify which voxels showed changes in that diffusion metric across groups while being agnostic to which conditions might differ. From these whole-brain ANOVAs, we next extracted the whole-brain t-stat maps that contrasted all pairs of treatment conditions for each diffusion metric and thresholded these at a voxelwise p<0.01. This threshold is not intended to identify reliable voxels or clusters that differentiate across groups but rather to provide an intentionally lenient initial threshold that would filter out the large number of voxels that showed little or no evidence of discrimination. Next, we merely counted the number of voxels that passed this threshold in either direction of the contrast. This whole-brain voxel count would be our outcome metric to analyze the effects of radiation on the pattern of diffusion metrics. To derive an estimate of the base rate, we performed a Monte Carlo analysis, randomly shuffling mice and assigning them to conditions, running the corresponding ANOVA, and extracting the resulting t-maps for a total of 10,000 randomized t-tests per metric. This allowed us to estimate the true alpha rate – the probability that the observed number of voxels crossing the voxelwise threshold of p<0.01 might be observed by chance.

Following this primary whole-brain analysis, we conducted an exploratory analysis to understand how widespread these effects might be. The analysis followed the methods used in the primary whole-brain analysis but focused on regions defined by the Allen Brain Atlas [22]. We counted the proportion of voxels crossing a threshold for each region within the atlas and compared this with estimates of the null derived from Monte Carlo simulations. Regions that exceeded p<0.05 versus that null were overlapped with averaged structural T2w TurboRare brains.

## Results

Ten months after irradiation, all mice were subjected to the NOR test. Only one outlier was removed according to the exclusion criteria. Whereas animals that received CONV-RT showed a decline in cognitive function, animals that received FLASH-RT performed similarly to the controls, suggesting cognitive protection (Fig. 2).

**Figure 2.**
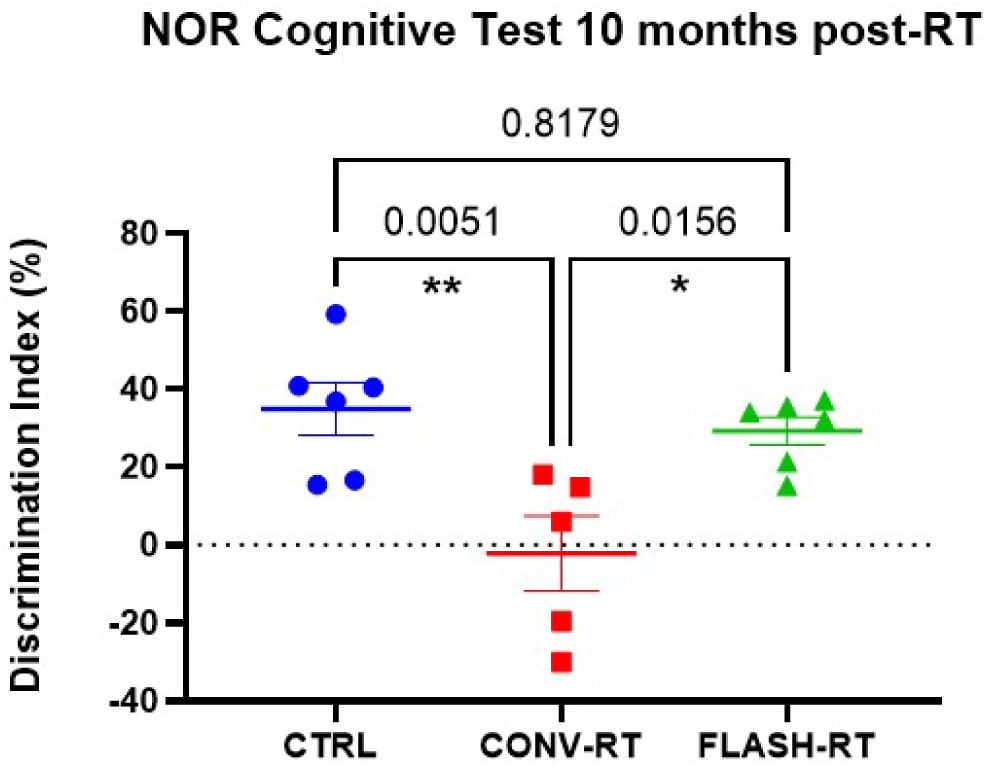
FLASH-RT spares long-term memory deficits. Scatter bar plots of the discrimination index (Eq.1) obtained after Novel Object Recognition cognitive testing ten months after treatment. Each discrimination data point represents one animal. P values plotted are the results of the Tukey post hoc test. Error bars represent SEM ( Standard Error of the Mean).

The volume and mean intensity of the hippocampus of each image were calculated, and the mean intensity was adjusted using Equation 2 above. According to the criteria, only two outliers were removed from the hippocampal volume analysis, and one outlier was removed for the mean intensity analysis. We observed some evidence for a decrease in hippocampal volume (8.3% average reduction; Fig 3A) in the CONV-RT cohort relative to the unirradiated cohort (Bonferroni test p=0.0621), but no difference was observed between the volume of the unirradiated cohort and the FLASH-RT cohort (3.3% average reduction; Bonferroni test p=0.9533; uncorrected one-way ANOVA p=0.0638, F(2,23)=3.108; Fig 3B). A statistically significant reduction in adjusted mean intensity of the hippocampus was observed between the unirradiated and CONV-RT animals (Bonferroni test p=0.0006) that was also not seen in the FLASH-RT ones (Bonferroni test p=0.1560; one-way ANOVA p=0.0008, F(2,24)=9.634; Fig 2C).

**Figure 3.**
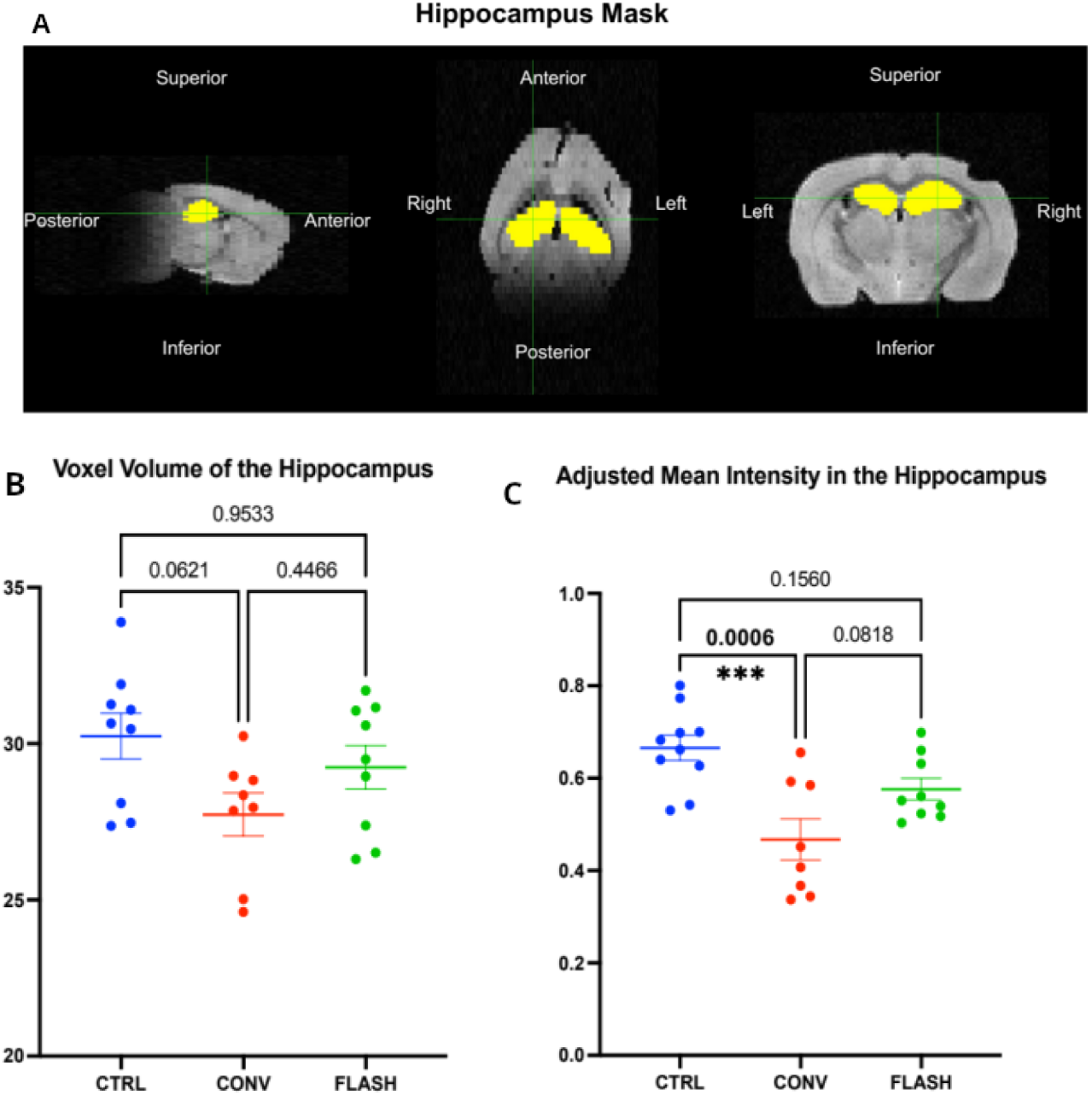
Hippocampal changes were observed in CONV-RT mice but not in FLASH-RT mice in T2 weighted structural MRI. A) The T2 weighted structural image of a control mouse with overlayed transformed masks of the hippocampus. B-C) Scatter bar plots of the volume and adjusted mean intensity of transformed masks. Each data point represents one animal. P values plotted are the results of the Bonferroni multiple comparisons test. Error bars represent SEM B) Total volume of the hippocampus mask in mm3. C) Mean Intensity of the hippocampus mask adjusted for the mean intensity of the whole brain using Equation 2.

Subsequent analysis was comparing the diffusion metrics across the three groups, as these metrics are designed to assess aspects of the microstructure of the tissue. As we had no *a priori* prediction concerning the location within the brain or direction of the change for each metric, our approach was to count the number of voxels that showed a difference between groups (defined as a p<0.01) and to compare this to the number that would be expected by chance (as estimated from a Monte Carlo permutation analysis). Results indicate directionality, but the statistics were performed without reference to direction unless otherwise indicated.

Comparing CONV-RT to Control, we found alterations in half of the diffusion metrics examined (Fig 4). The AD, ADC, RD, and QIV metrics all showed several voxels with decreases in their value following CONV-RT and few, if any, exhibiting increases (QIV p <0.05, all other p’s <0.01). In contrast, RTAP, RTOP, and RTPP showed the converse, increasing their value following CONV-RT (RTOP p<0.01, all others p<0.05). Averaging across all 14 metrics, 4746 voxels, which represents 2.7% of the brain, showed a change following CONV-RT (p<0.0001).

**Figure 4.**
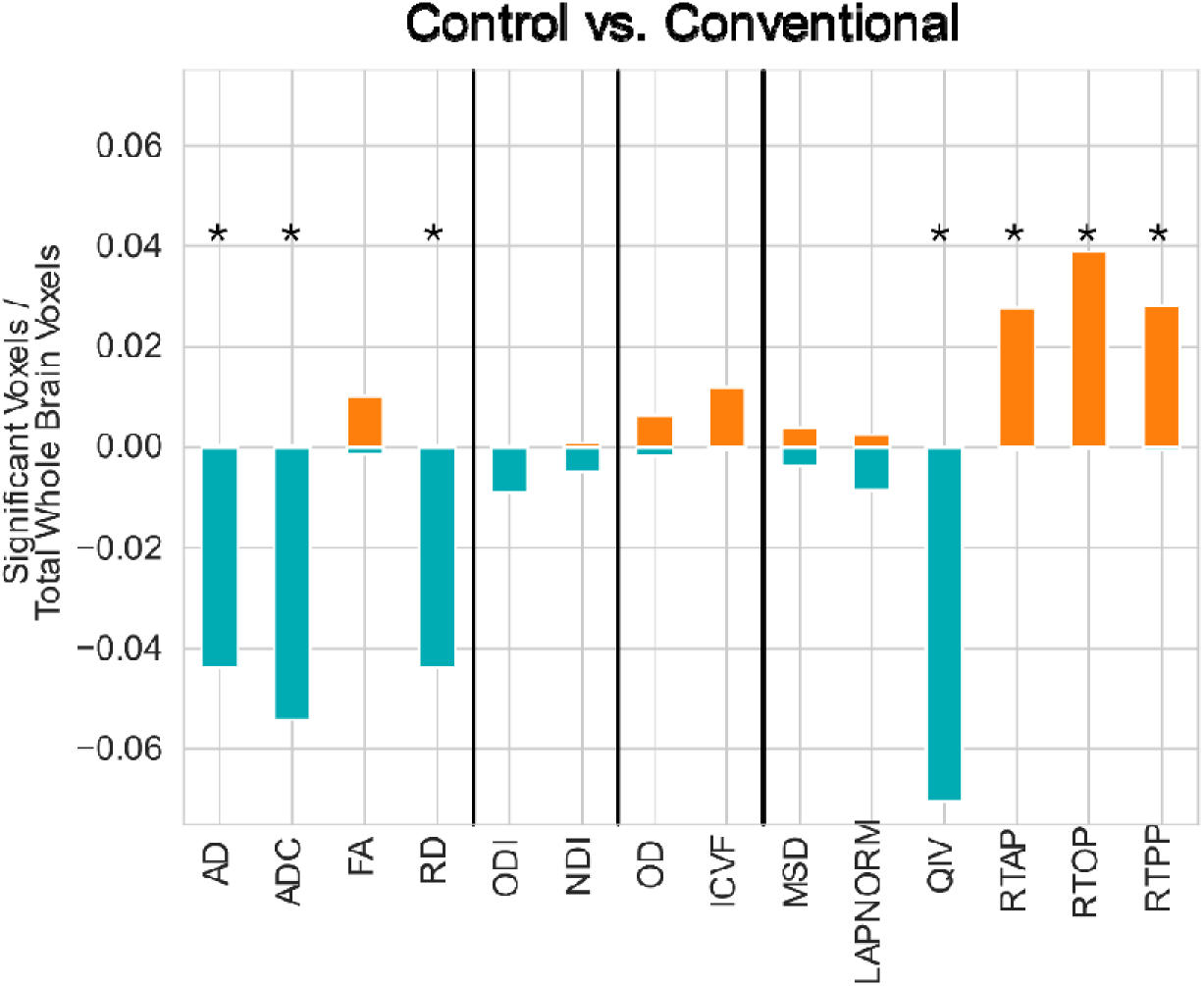
CONV-RT substantially altered brain diffusion measures relative to Control. Percentage of voxels showing increase (orange) or decrease (blue) in diffusion metrics post-conventional irradiation vs control. Collapsing across the brain, the number of voxels exhibiting treatment-induced changes in diffusion values (combined height of orange and blue bars) far exceeded that expected by chance, with reliable increases and decreases observed in every metric.

In contrast, FLASH had minimal impact on diffusion within the brain (Fig 5A). While the pattern of increases and decreases followed that of CONV-RT, it was markedly reduced. No metrics showed any significant change from control (all p’s > 0.17).

**Figure 5.**
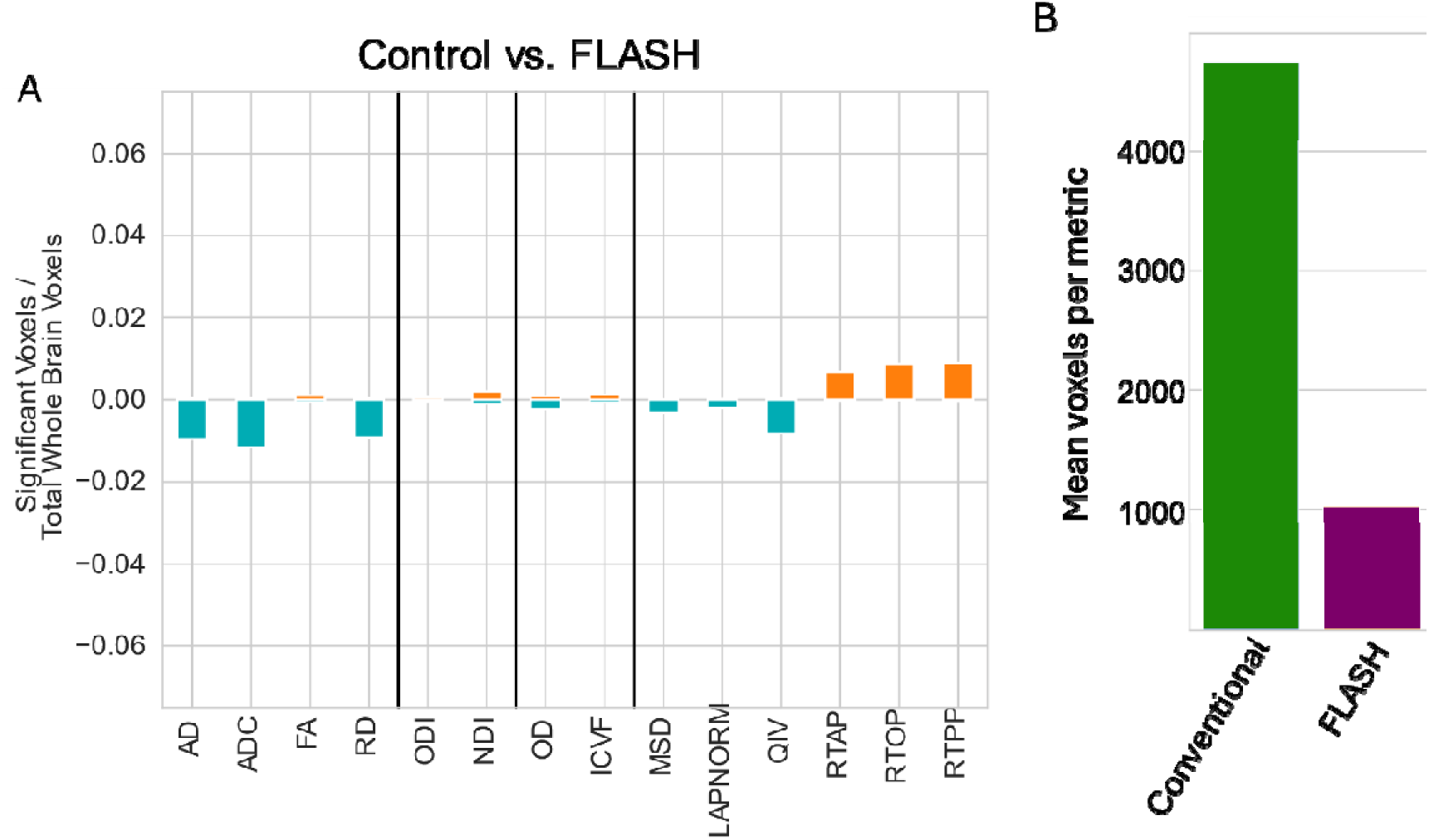
Control vs FLASH-RT showed a similar but heavily attenuated pattern in brain diffusion. A) Percentage of voxels showing increase (orange) or decrease (blue) in diffusion metrics post-conventional irradiation vs control. Collapsing across the brain, the number of voxels exhibiting treatment-induced changes in diffusion values (combined height of orange and blue bars) followed the same pattern as Conventional irradiation but with a markedly attenuated effect. B) While reliable effects were observed in a number of the metrics, they were found in only 21% as many voxels as Conventional.

Averaged across all metrics, only 1029 voxels were affected by FLASH-RT (0.58% of the brain). Thus 4.6x as many voxels were affected by CONV-RT irradiation as FLASH-RT (Fig 5B). Then, the probability of each metric was compared against the null distribution. The comparison between Control and CONV-RT, showed that some metrics displayed smaller p-values than the null distribution, indicating a more substantial deviation to the right (Figure 6). Conversely, the comparison between Control and FLASH-RT showed an alignment close to the null distribution. When FLASH-RT *versus* CONV-RT were compared, some metrics exhibited lower p-values than those observed in the Control *versus* FLASH comparison, but the effects were not significant. This indicates that the effect of CONV-RT but not FLASH-RT relative to Control is robust for specific metrics, while FLASH-RT response is more intermediate.

**Figure 6.**
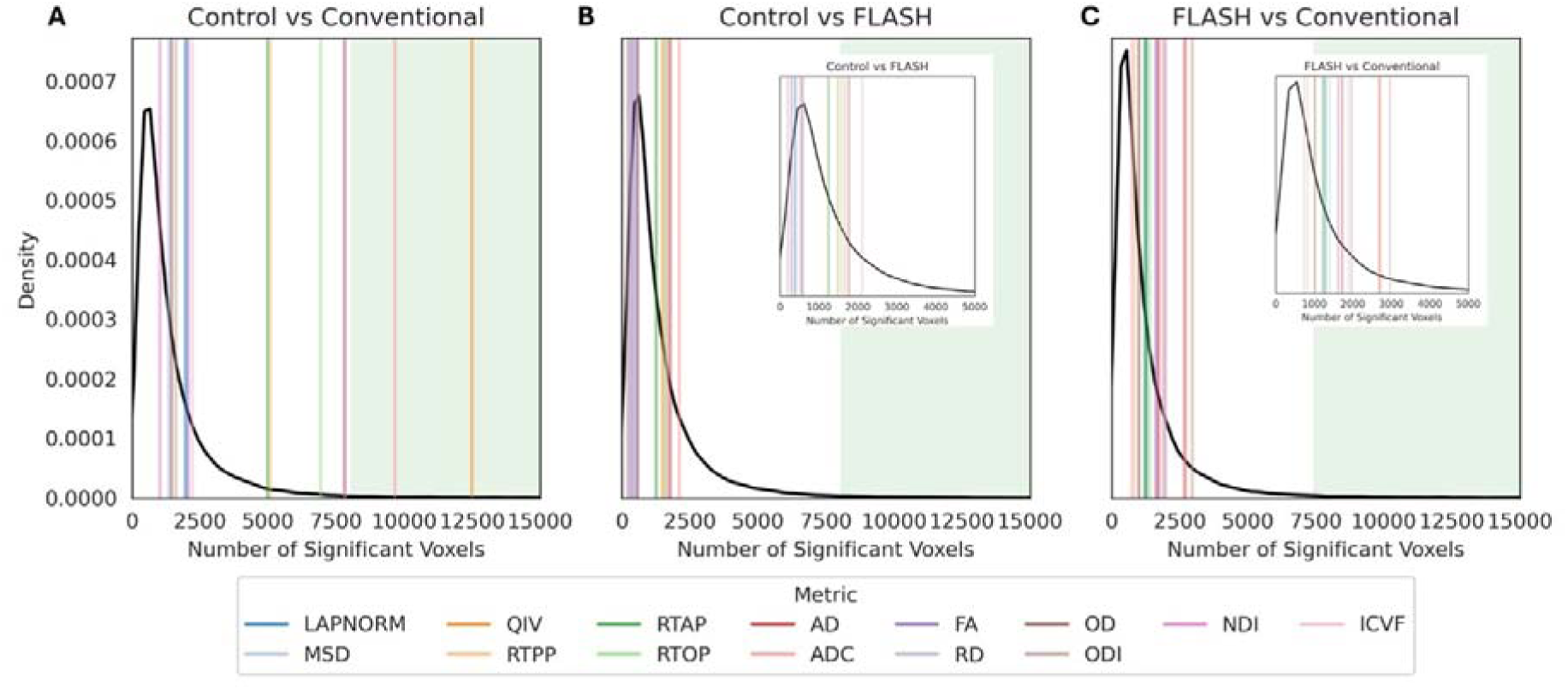
After CONV-RT, the p values are smaller than for the Null distribution for several metrics, but not after FLASH-RT. In each panel, the black line represents the null distribution, while the vertical lines indicate the observed data for each diffusion metric. The p-value corresponds to the area under the tail to the right of each metric’s vertical line. p-values in the green zone are significant at the.01 level. A) CONV-RT vs Controls, B) FLASH-RT vs Controls C) FLASH-RT vs CONV-RT

Consistently with the results collected across the whole brain (Figure 5B), in Figures 7 and 8, a decrease in AD, ADC, QIV, and RD is associated with CONV-RT that were markedly reduced in FLASH-RT (Figure 7), with most prominent decreases found in the medial septal nucleus, the globus pallidus, insular cortex, primary somatosensory area, and the parastrial nucleus (Supplementary Tables S1-S4). Increases in RTAP, RTOP, and RTPP metrics are associated with CONV-RT, and are reduced in FLASH-RT (Figure 8), most prominently in the same regions (Supplementary Tables S5-S7). The complete list of regions with changes is presented in Supplemental Tables S1-S7.

**Figure 7.**
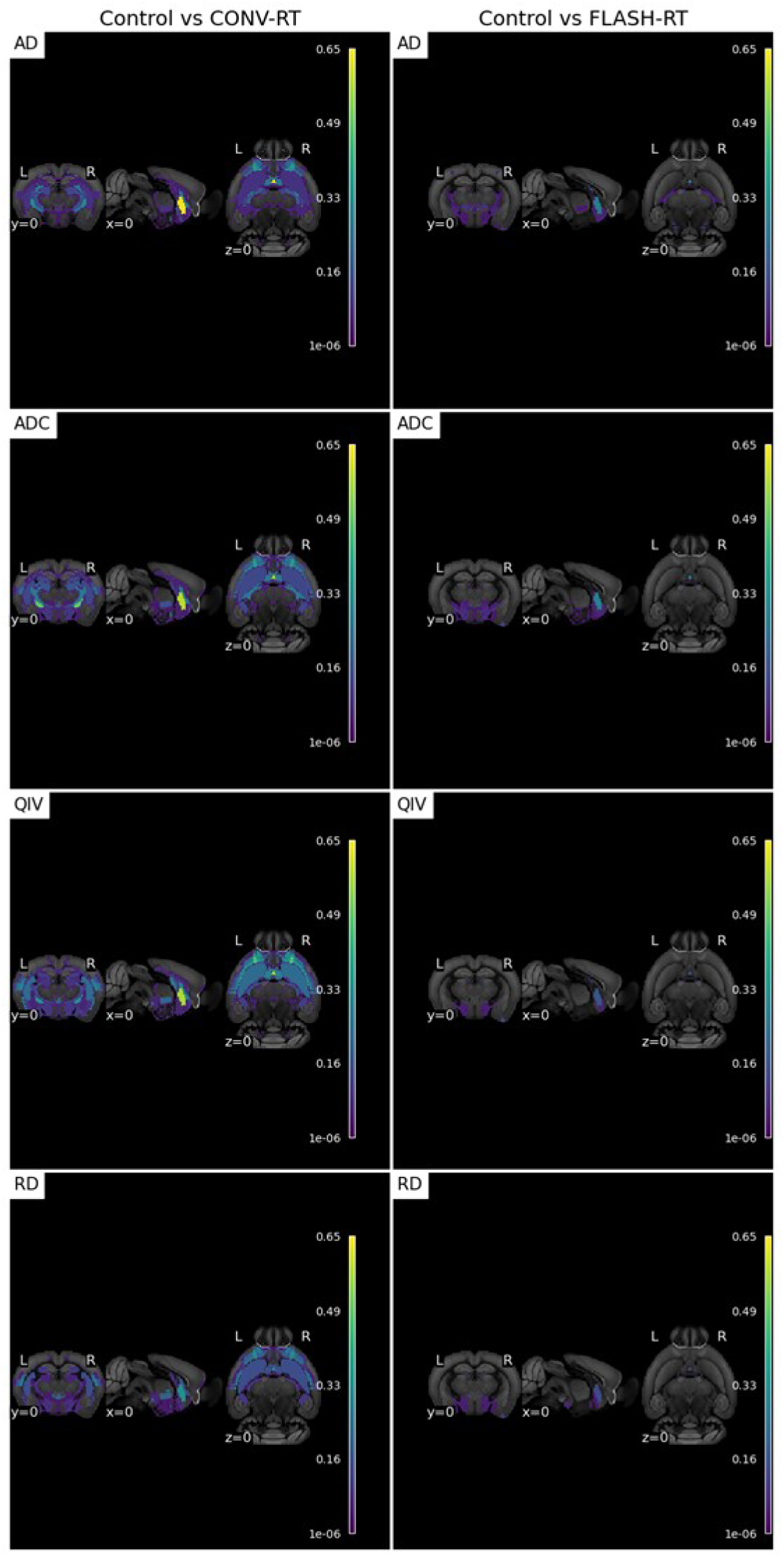
Regional changes in diffusion in the brain following CONV-RT relative to Controls (left) and FLASH-RT relative to Controls (right) in AD, ADC, QIV, and RD diffusion metrics. Color code represents the proportion of voxels crossing a threshold for each region within the atlas. Voxels were compared with estimates of the null derived from Monte Carlo simulations and included only when exceeded p<0.05 versus that null. Color code is overlapped with averaged structural T2w TurboRare brains.

**Figure 8.**
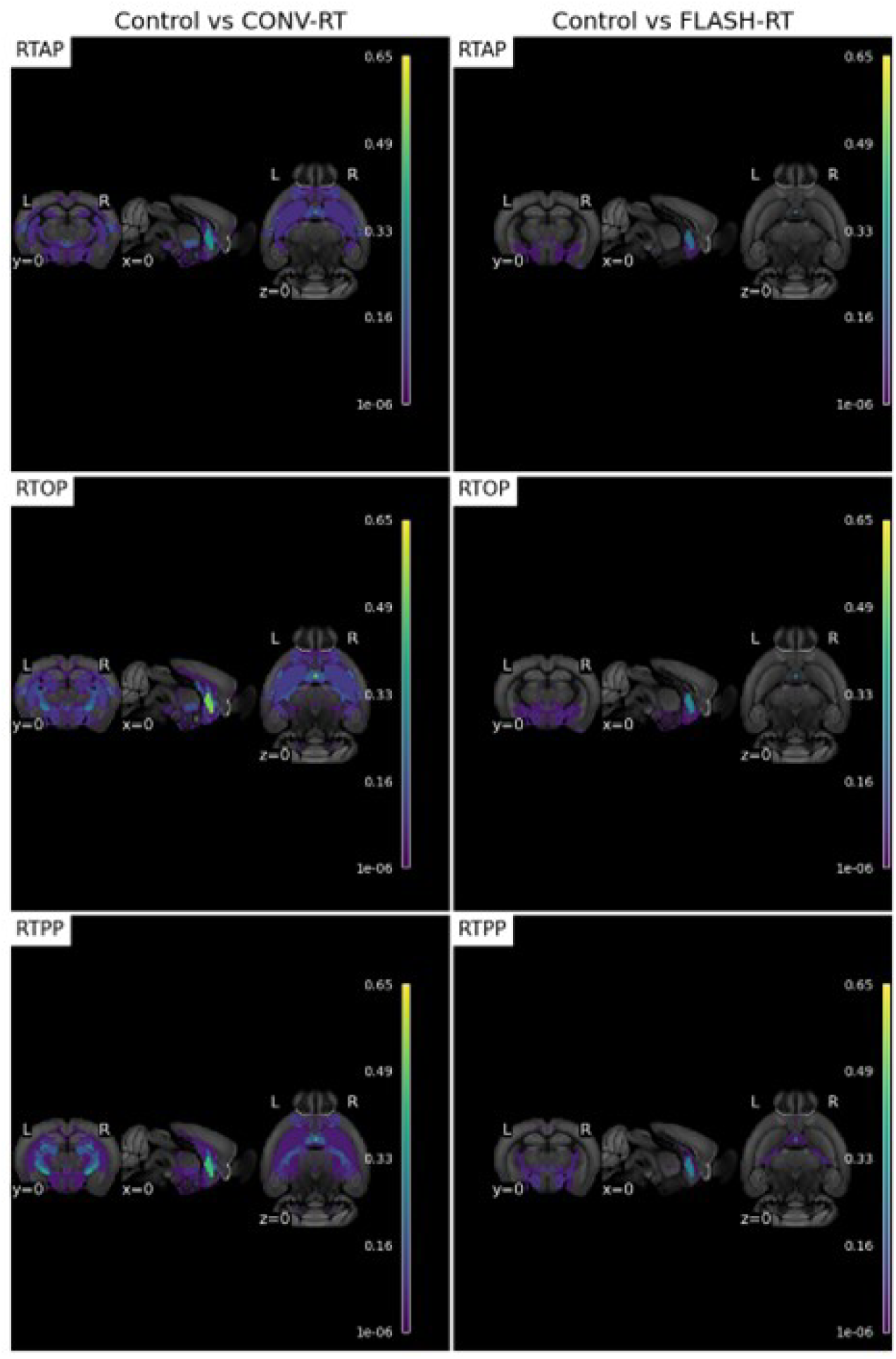
Regional changes in diffusion in the brain following CONV-RT relative to Controls (left) and FLASH-RT relative to Controls (right) in RTAP, RTOP, and RTPP diffusion metrics. Color code represents the proportion of voxels crossing a threshold for each region within the atlas. Voxels were compared with estimates of the null derived from Monte Carlo simulations and included only when exceeded p<0.05 versus that null. Color code is overlapped with averaged structural T2w TurboRare brains.

## Discussion

In the present study, we showed that ex-vivo structural and diffusion-weighted imaging (DWI) MRI can reveal changes induced by CONV-RT versus non-irradiated controls at a late post-irradiation time, whereas no significant modifications were found between FLASH-RT and controls. However, differences between CONV-RT and FLASH-RT remained non-statistically different, which could be influenced by the sample size but also suggest that FLASH-RT still induces intermediate structural and microstructural damage to the brain.

Using the same cohort of animals, cognition evaluated using the NOR test was preserved after exposure to FLASH-RT but not CONV-RT. These findings corroborate many cognitive studies published in the past after FLASH-RT [1, 2, 3, 4, 18, 37, 38]. Data also indicate that the performance of older control animals is inferior to that of younger animals [1]. However, no linear correlation was found between MRI and cognitive results, highlighting the difficulty of linking cognitive deficits to specific MRI signatures.

When T2-weighted MRIs were analyzed, the hippocampal volume showed a decrease of 8.3% after CONV-RT compared to controls, whereas after FLASH-RT, the decrease was only 3.3%. The reduction remained non-statistically significant, but the trend could be noteworthy as decreased hippocampal volume has been linked to cognitive decline in animal models of diseases like Alzheimer’s (AD), Parkinson’s, and Huntington’s [8, 39, 40]. Similarly, standard radiotherapy has also been shown to decrease the hippocampal volume with a mean reduction of 5.7%, 4-7 months post-radiotherapy [41].

When analysis on hippocampal T2-signal intensity was performed, a statistically significant reduction was found after CONV-RT compared to controls but not after FLASH-RT. These differences in intensity against controls could mean that CONV-RT but not FLASH-RT could decrease the free water content in hippocampal tissues. This might reflect regional tissue damage and/or loss of tissue integrity similar to the effects observed during aging or AD [11, 12, 42, 43]. These results further suggest that preservation of hippocampal structure following FLASH-RT, as monitored by MRI, could, at least in part, contribute to cognitive protection along with previously found reduced neuroinflammation and preservation of the microvasculature [1, 2, 18, 38, 37]. Again, when the T2-signal intensity after CONV-RT vs. FLASH-RT was compared, the observed mean intensity was lower in the CONV-RT versus the FLASH-RT group, but the difference was not statistically significant (p=.0818).

Since recent advancements in diffusion-weighted imaging (DWI) techniques have been shown to detect structural and microstructural changes after cranial radiation, we performed a similar analysis on our cohort of animals. Clinical studies performed after standard dose rate irradiation have shown changes in traditional diffusion metrics suggestive of dose-dependent demyelination and axonal injury [44, 45]. Similar results have been found in preclinical models (porcine, rat, mouse), where white matter was more susceptible to necrosis than grey matter, each exhibiting distinct dose tolerances [46, 47, 48]. Additionally, DWI analysis has been shown to correlate with cognitive deficits observed in some rodent models [49]. Our results are consistent with these published results, as significant changes in DWI were measured after CONV-RT. The reported diffusion metrics exhibit significant variations from the values expected by chance across the entire brain. Interestingly, no changes in DWI metrics were measured after FLASH-RT. Translating MRI signatures to specific biological substrates and processes involved in the progression and manifestation of neurocognitive decline has proven challenging. Consistently, in our study, according to Spearman’s rank test, we found no statistically significant correlations between the individual discrimination index values and individual structural MRI changes. Similarly, for DWI metrics, the linear model did not find a direct relationship with the discrimination index. The limited sample size included in our study (n=6 animals) likely contributed to this lack of correlation. However, the absence of a direct relationship could also suggest the presence of distinct underlying factors influencing diffusion outcomes and behavior separately or might reflect that complex regional relationships can not be monitored using our whole-brain analysis strategy, which is inherently less sensitive. In efforts to address the issues inherent to small sample sizes, the statistical approach chosen to analyze the whole brain DTI images specifically considers such limitations, as detailed in the methods section. To enhance sensitivity while maintaining reliability, our primary analysis was conducted across the entire brain and made no attempt to identify the location of any effects. This choice was made as a proof of principle study using an ex-vivo imaging strategy that reduces signal-to-noise and motion artifacts. Additionally, DTI analysis using ex vivo brains has previously shown to be highly sensitive for detecting white matter damage, despite changes in diffusivity parameters when compared to in vivo analyses [50].

In an attempt to identify certain effects in specific regions of the brain, we conducted an exploratory analysis examining each ROI independently (without correcting for multiple comparisons across ROIs). According to this exploratory regional analysis (Supplementary Tables S1-S7), the basal forebrain (specifically, the Medial Septal Nucleus, MSN) and the insular cortex show changes in almost all diffusion metrics after CONV-RT. Interestingly, the MSN metrics were unaffected or showed minimal changes after FLASH-RT. This is important because the MSN sends cholinergic, GABAergic, and glutamatergic projections to the hippocampus, constituting a crucial circuit for generating and modulating hippocampal theta rhythms associated with attention, learning, and memory consolidation [51]. Therefore, the minimal changes found after FLASH-RT could be linked to the observed memory protection found on the NOR cognitive test. Similarly, changes in diffusion metrics were observed in the insular cortex after CONV-RT but not FLASH-RT. As the agranular insular cortex is related to integrating emotional, sensory, and cognitive information, it could also positively influence the NOR test results obtained after FLASH-RT. The paraventricular hypothalamic nucleus also showed modified diffusion metrics after CONV-RT but not FLASH-RT. The paraventricular hypothalamic nucleus is a key regulator of stress responses and circadian rhythms, including sleep [52]. CONV-RT is known to produce sleep disturbances in patients, referred to as radiation somnolence syndrome [53], and in rodent models, it is known to increase sleep during the active phase [54] and modify hypothalamic levels of neurotransmitters [55] and neuroinflammation markers [54]. Therefore, the observed changes in hypothalamic diffusion metrics could also be related to this known hypothalamic radiation response, which might be attenuated after FLASH-RT. Interestingly, two additional diffusion metrics were shown to be modified in the parastrial nucleus of the preoptic area of the hypothalamus, but only in FLASH-RT treated animals. Since the parastrial nucleus is involved in homeostasis and critical for regulating emotional and motivational states and sleep it might be associated with better outcomes after FLASH-RT. Clearly, further studies are needed to confirm whether FLASH-RT protects hypothalamic homeostasis compared to CONV-RT.

In conclusion, this study is the first to show that FLASH-RT as compared with controls does not induce any changes detectable by MRI at long-term, whereas structural and microstructural alterations are detected after CONV-RT compared to controls. While no differential effect between FLASH-RT and CONV-RT was measured, this study suggests that more investigations should be performed using MRI to investigate the sparing effect of FLASH-RT on the normal brain.

## Acknowledgments

Funding was provided by National Institutes of Health grant P01CA244091-01 (to CLL & MCV supporting JJ & VG); Swiss National Science Foundation grantSpirit IZSTZ0_198747/1 (to MCV and PBZ supporting JFC) and MAGIC-FNS CRSII5_186369 (to MCV and supporting VG); and CONACYT for supporting PBZ sabbatical in Switzerland. We acknowledge the resources and expertise of the CIBM Center for Biomedical Imaging. The authors thank Dr. Cristina Cudalbu for all her helpful comments, Dr. Yohan Van De Looij for insightful advice on the sample setup, and Dr. Katarzyna Pierzchala for her help with the sample preparation. We would also like to express our gratitude to Dr. Analina Housin and Stefanita-Octavian Mitrea for their support with brain extraction procedures. Finally, we would like to thank the core facilities and staff at Lausanne CHUV, as well as their veterinary staff, for their valuable support.

## Disclosure

No disclosures to declare.

## Declaration of generative AI and AI-assisted technologies in the writing process

During the preparation of this work the author(s) used Grammarly in order to check the English. After using this tool/service, the author(s) reviewed and edited the content as needed and take(s) full responsibility for the content of the publication.

## Notes

### Competing Interest Statement

The authors have declared no competing interest.

