## Supplemental Tables for "Differentiating unirradiated mice from those exposed to conventional or FLASH radiotherapy using MRI"

**Supplementary Data**

Table S1. Ratio of AD Decreasing Voxels / Region Size by Region and Comparison

| ROI ID | Region Name | Comparison | Ratio |
| --- | --- | --- | --- |
| 564 | Medial septal nucleus | Control vs CONV-RT | 0.636 |
| 800 | Agranular insular area ventral part layer 5 | Control vs CONV-RT | 0.380 |
| 440 | Orbital area lateral part layer 6a | Control vs CONV-RT | 0.360 |
| 470 | Subthalamic nucleus | Control vs CONV-RT | 0.340 |
| 783 | Agranular insular area dorsal part layer 6a | Control vs CONV-RT | 0.336 |
| 30 | Periventricular hypothalamic nucleus anterior part | Control vs CONV-RT | 0.333 |
| 675 | Agranular insular area ventral part layer 6a | Control vs CONV-RT | 0.333 |
| 1101 | Agranular insular area dorsal part layer 5 | Control vs CONV-RT | 0.327 |
| 1022 | Globus pallidus external segment | Control vs CONV-RT | 0.321 |
| 363 | Prelimbic area layer 5 | Control vs CONV-RT | 0.285 |

Table S2. Ratio of ADC Decreasing Voxels / Region Size by Region and Comparison

| ROI ID | Region Name | Comparison | Ratio |
| --- | --- | --- | --- |
| 564 | Medial septal nucleus | Control vs CONV-RT | 0.580 |
| 1031 | Globus pallidus internal segment | Control vs CONV-RT | 0.498 |
| 929 | Primary somatosensory area nose layer 6b | Control vs CONV-RT | 0.491 |
| 1109 | Parastrial nucleus | Control vs FLASH-RT | 0.417 |
| 30 | Periventricular hypothalamic nucleus anterior part | Control vs CONV-RT | 0.417 |
| 783 | Agranular insular area dorsal part layer 6a | Control vs CONV-RT | 0.413 |
| 800 | Agranular insular area ventral part layer 5 | Control vs CONV-RT | 0.385 |
| 675 | Agranular insular area ventral part layer 6a | Control vs CONV-RT | 0.375 |
| 1101 | Agranular insular area dorsal part layer 5 | Control vs CONV-RT | 0.340 |
| 849 | Visceral area layer 6b | Control vs CONV-RT | 0.308 |

Table S3. Ratio of QIV Decreasing Voxels / Region Size by Region and Comparison

| ROI ID | Region Name | Comparison | Ratio |
| --- | --- | --- | --- |
| 564 | Medial septal nucleus | Control vs CONV-RT | 0.567 |
| 783 | Agranular insular area dorsal part layer 6a | Control vs CONV-RT | 0.559 |
| 675 | Agranular insular area ventral part layer 6a | Control vs CONV-RT | 0.521 |
| 800 | Agranular insular area ventral part layer 5 | Control vs CONV-RT | 0.498 |
| 893 | Supplemental somatosensory area layer 6b | Control vs CONV-RT | 0.409 |
| 1101 | Agranular insular area dorsal part layer 5 | Control vs CONV-RT | 0.369 |
| 341 | supramammillary decussation | Control vs CONV-RT | 0.357 |
| 884 | amygdalar capsule | Control vs CONV-RT | 0.351 |
| 849 | Visceral area layer 6b | Control vs CONV-RT | 0.346 |
| 1125 | Orbital area ventrolateral part layer 5 | Control vs CONV-RT | 0.344 |

Table S4. Ratio of RD Decreasing Voxels / Region Size by Region and Comparison

| ROI ID | Region Name | Comparison | Ratio |
| --- | --- | --- | --- |
| 1109 | Parastrial nucleus | Control vs FLASH-RT | 0.417 |
| 783 | Agranular insular area dorsal part layer 6a | Control vs CONV-RT | 0.385 |
| 800 | Agranular insular area ventral part layer 5 | Control vs CONV-RT | 0.352 |
| 564 | Medial septal nucleus | Control vs CONV-RT | 0.329 |
| 1101 | Agranular insular area dorsal part layer 5 | Control vs CONV-RT | 0.298 |
| 675 | Agranular insular area ventral part layer 6a | Control vs CONV-RT | 0.292 |
| 341 | supramammillary decussation | Control vs CONV-RT | 0.286 |
| 893 | Supplemental somatosensory area layer 6b | Control vs CONV-RT | 0.273 |
| 849 | Visceral area layer 6b | Control vs CONV-RT | 0.269 |
| 520 | Ventral auditory area layer 6a | Control vs CONV-RT | 0.261 |

Table S5. Ratio of RTAP Increasing Voxels / Region Size by Region and Comparison

| ROI ID | Region Name | Comparison | Ratio |
| --- | --- | --- | --- |
| 564 | Medial septal nucleus | Control vs CONV-RT | 0.326 |
| 30 | Periventricular hypothalamic nucleus anterior part | Control vs CONV-RT | 0.250 |
| 849 | Visceral area layer 6b | Control vs CONV-RT | 0.231 |
| 564 | Medial septal nucleus | Control vs FLASH-RT | 0.226 |
| 893 | Supplemental somatosensory area layer 6b | Control vs CONV-RT | 0.197 |
| 694 | Agranular insular area ventral part layer 2 3 | Control vs CONV-RT | 0.196 |
| 520 | Ventral auditory area layer 6a | Control vs CONV-RT | 0.187 |
| 181 | Nucleus of reunions | Control vs CONV-RT | 0.177 |
| 1077 | Perireunensis nucleus | Control vs CONV-RT | 0.167 |
| 800 | Agranular insular area ventral part layer 5 | Control vs CONV-RT | 0.164 |

Table S6. Ratio of RTOP Increasing Voxels / Region Size by Region and Comparison

| ROI ID | Region Name | Comparison | Ratio |
| --- | --- | --- | --- |
| 564 | Medial septal nucleus | Control vs CONV-RT | 0.517 |
| 30 | Periventricular hypothalamic nucleus anterior part | Control vs CONV-RT | 0.500 |
| 929 | Primary somatosensory area nose layer 6b | Control vs CONV-RT | 0.396 |
| 564 | Medial septal nucleus | Control vs FLASH-RT | 0.301 |
| 1031 | Globus pallidus internal segment | Control vs CONV-RT | 0.295 |
| 849 | Visceral area layer 6b | Control vs CONV-RT | 0.269 |
| 287 | Bed nucleus of the anterior commissure | Control vs CONV-RT | 0.250 |
| 66 | Lateral terminal nucleus of the accessory optic tract | Control vs CONV-RT | 0.250 |
| 1102 | Primary somatosensory area mouth layer 6a | Control vs CONV-RT | 0.233 |
| 694 | Agranular insular area ventral part layer 2 3 | Control vs CONV-RT | 0.221 |

Table S7. Ratio of RTPP Increasing Voxels / Region Size by Region and Comparison

| ROI ID | Region Name | Comparison | Ratio |
| --- | --- | --- | --- |
| 564 | Medial septal nucleus | Control vs CONV-RT | 0.442 |
| 1031 | Globus pallidus internal segment | Control vs CONV-RT | 0.391 |
| 470 | Subthalamic nucleus | Control vs CONV-RT | 0.381 |
| 603 | fimbria | Control vs CONV-RT | 0.277 |
| 564 | Medial septal nucleus | Control vs FLASH-RT | 0.273 |
| 363 | Prelimbic area layer 5 | Control vs CONV-RT | 0.271 |
| 910 | Orbital area medial part layer 6a | Control vs CONV-RT | 0.266 |
| 1022 | Globus pallidus external segment | Control vs CONV-RT | 0.264 |
| 66 | Lateral terminal nucleus of the accessory optic tract | Control vs CONV-RT | 0.250 |
| 250 | Lateral septal nucleus caudal (caudodorsal) part | Control vs CONV-RT | 0.197 |
